# Intrinsic limitations in mainstream methods of identifying network motifs in biology

**DOI:** 10.1101/272401

**Authors:** James Fodor, Michael Brand, Rebecca J Stones, Ashley M Buckle

## Abstract

Network motifs are connectivity structures that occur with significantly higher frequency than chance, and are thought to play important roles in complex biological networks, for example in gene regulation, interactomes, and metabolomes. Network motifs may also become pivotal in the rational design and engineering of complex biological systems underpinning the field of synthetic biology. Distinguishing true motifs from arbitrary substructures, however, remains a challenge. Here we demonstrate both theoretically and empirically that implicit assumptions present in mainstream methods for motif identification do not necessarily hold, with the ramification that motif studies using these mainstream methods are less able to effectively differentiate between spurious results and events of true statistical significance than is often presented. We show that these difficulties cannot be overcome without revising the methods of statistical analysis used to identify motifs. The implications of these findings are therefore far-reaching across diverse areas of biology.

---

A *network motif* is a particular connected pattern of nodes and edges that appears in a network significantly more frequently than would be expected by chance. It is hypothesized that network motifs play a more important role in network function than arbitrary substructures. Since the concept was popularised in 2002 by Milo et al. [1, 2], much effort has been devoted to identifying network motifs in the hope that doing so will yield insights into network behaviour [3-5]. Network motif identification plays an important role in molecular and cell biology research, notably the study of gene regulation [3] (which describes regulatory relationships between transcription factors and their target genes), interactomes [6] (which describe protein-protein interactions), and metabolomes [7] (which describe the complete set of small molecules within a cell).

In the study of network motifs, it is critical to be able to determine whether a particular subgraph *H* (i.e. a specific connected pattern of nodes and edges) observed in a network *G* is in fact a motif, or merely a chance occurrence. In order to determine whether *H* is a motif of *G,* the following procedure (or a close variant thereof) is typically followed [8]:

- *Step 1*: Define 𝕊(*G*) to be the set of all networks *similar* to *G,* in the sense that they have the same set of vertices, each with the same number of incoming, outgoing and bi-directional edges.
- *Step 2:* Count the number *n*_*G*_*(H)* of times that the putative motif *H* occurs in *G.*
- *Step 3:* Estimate the probability that a network selected uniformly at random from the set 𝕊(*G*) will contain at least *n*_*G*_*(H)* copies of *H,* typically by use of an edge-switching algorithm [9]. (See Supplementary Information (SI) S4.4 for more information on edge-switching algorithms.)
- *Step 4:* If the estimated probability of at least *n*_*G*_*(H)* copies of *H* occurring by chance is less than some user-defined threshold, declare *H* to be a motif of *G.*

While the precise algorithms used to implement these steps differ between implementations, this underlying methodology is adopted by popular network motif software tools such as FANMOD [10], Mfinder [8], and MAVisto [11], and is widely used for identifying network motifs in a diverse range of biological systems. For example, Zhao et al. [12] identified twelve types of three-node motifs in chromatin-state transcriptional regulatory networks in four human cell lines. They first generated a set of 500 similar networks 𝕊(*G*) using the FANMOD tool, and then declared a candidate to be a motif if its frequency in the original network *G* was significantly greater than in the set of similar networks, using a significance threshold (p-value) of 0.05. We note that these are twelve identified motifs out of only thirteen motif candidates of that size in total. FANMOD was also used in identifying four types of three-node motifs in a miRNA-transcription factor (TF) co-regulatory network in non-small cell lung cancer, in this case with a candidate declared to be a motif on the basis of a 0.01 *p*-value threshold [13]. In a third example, Vinayagam et al. [14] analysed a protein-protein interaction network of *Drosophila melanogaster,* finding seven types of 3-node motifs using FANMOD to generate random sets of 1000 similar networks, with the significance of motifs evaluated using Z-scores.

In order to explore the assumptions inherent in mainstream methods of network motif detection, we analysed three real-world networks; the *E. coli* TF network, the *S. cerevisiae* TF network and the *E. coli* metabolic network. Both organisms are the best known and characterized genome model organisms. The mainstream methods outlined above were used in the analysis of both TF [15, 16] and metabolic networks [17, 18]. Drawing upon these empirical analyses and additional theoretical results, the following two sections discuss two assumptions that are inherent in practical implementations of this approach, namely the assumption of a normal distribution and the assumption of independence between candidates. We show that both these assumptions fail to hold in all cases, and that this has serious ramifications for present-day mainstream motif identification techniques. The final section discusses the problem of failing to properly distinguish motif frequency from concentration, and the implications of this distinction for motif detection methods.

## The normality assumption

Motif-searching involves testing a very large number of hypotheses in parallel, with an associated high risk of false positive results. For example, the total number of possible 6-node motif candidates is over 1.5 million, meaning that if all are tested for, roughly one such candidate motif will be expected to receive a *p*-value of *10*^-6^ or less purely by chance. If meaningful results are to be obtained, it is therefore critical to be able to reliably differentiate between very small *p*-values, such as between a *p*-value of 10^−6^ (likely to be a false positive in this example) and 10^−10^ (likely to be a true positive in this example). Reliably distinguishing between such small *p*-values based on empirical frequencies alone unfortunately often requires a sample size that is prohibitively large; for instance, a sample size on the order of ten billion would be required for events with *p*-values at the 10^−10^ level to be sampled at all. As such, in order to directly evaluate the probability that a given motif candidate will occur in a network by chance, it would be necessary to generate a much larger set of similar networks from 𝕊(*G*) than is practical using current methods. Without accurate calculation of such probabilities, however, steps 3 and 4 of the mainstream approach to motif detection cannot be performed. The customary solution to this problem has been to assume that the motif frequency *n*_*G*_*(H)* is normally distributed, thereby enabling *p*-values to be easily calculated from the corresponding Z-score, which does not require unrealistically large samples [19].

As Picard et al. have observed, however, the normal distribution can be a poor approximation of the true motif distribution [20] (See also SI section S2.6 for other previous criticisms of motif detection methods). To illustrate this phenomenon, in Figure 1 we explicitly identify three synthetic examples of networks and associated motif candidates that follow three completely different distributions: Poisson, binomial, and a multimodal distribution. (Detailed explanations of the example networks and proofs of the stated asymptotic distributions appear in SI sections S5.2-S5.4, and also figure S1.) Strikingly, in this example a single subgraph follows different distributions in different networks, with the same subgraph following a binomial distribution in one class of networks (middle column) and a multimodal distribution in a different class of networks (right column). In such circumstances, assuming a normal distribution will lead to misleading *p*-values, which in this case will be highly significance-inflated.

**Figure 1:**
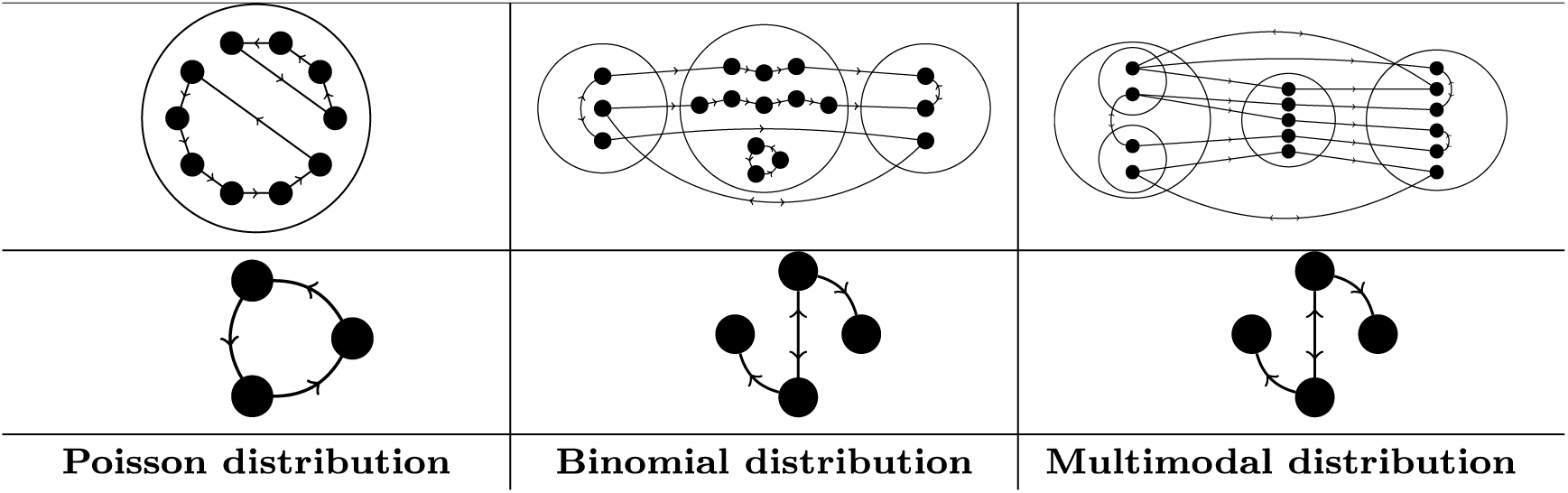
Three examples of classes of synthetic networks and corresponding motifs that exhibit non-normal asymptotic frequency distributions. Top row; An illustrative network from each class. Middle row; A motif candidate. Bottom row; The form of the frequency distribution of the given motif candidate, calculated for networks drawn from the corresponding class.

Furthermore, the usual definition of the *p*-value (the probability under the null hypothesis of obtaining results at least as extreme as what is actually observed) implicitly assumes that more extreme results are correspondingly less likely to occur. While this holds for the normal distribution, for severely multimodal distributions more extreme results can actually be *more* likely to occur than less extreme results, meaning that in such cases *p*-values calculated from Z-scores have no clear statistical interpretation (for an illustration see SI Figure S2).

Given the variety of behaviours we observe in these simple examples, we hypothesize that real-world networks feature an unpredictable hodgepodge of behaviours governing the distribution of subgraph frequencies in a set of similar networks. Analysis of *E. coli* TF and yeast networks indeed shows this to be the case (Figure 2; see also SI Figure S3 for more examples). While these examples are all clearly non-normal over the displayed domain, the synthetic examples from Figure 1 complement these real-world data by showing that such non-normality can continue far into the tail of the distribution, beyond those regions which can be easily sampled.

**Figure 2:**
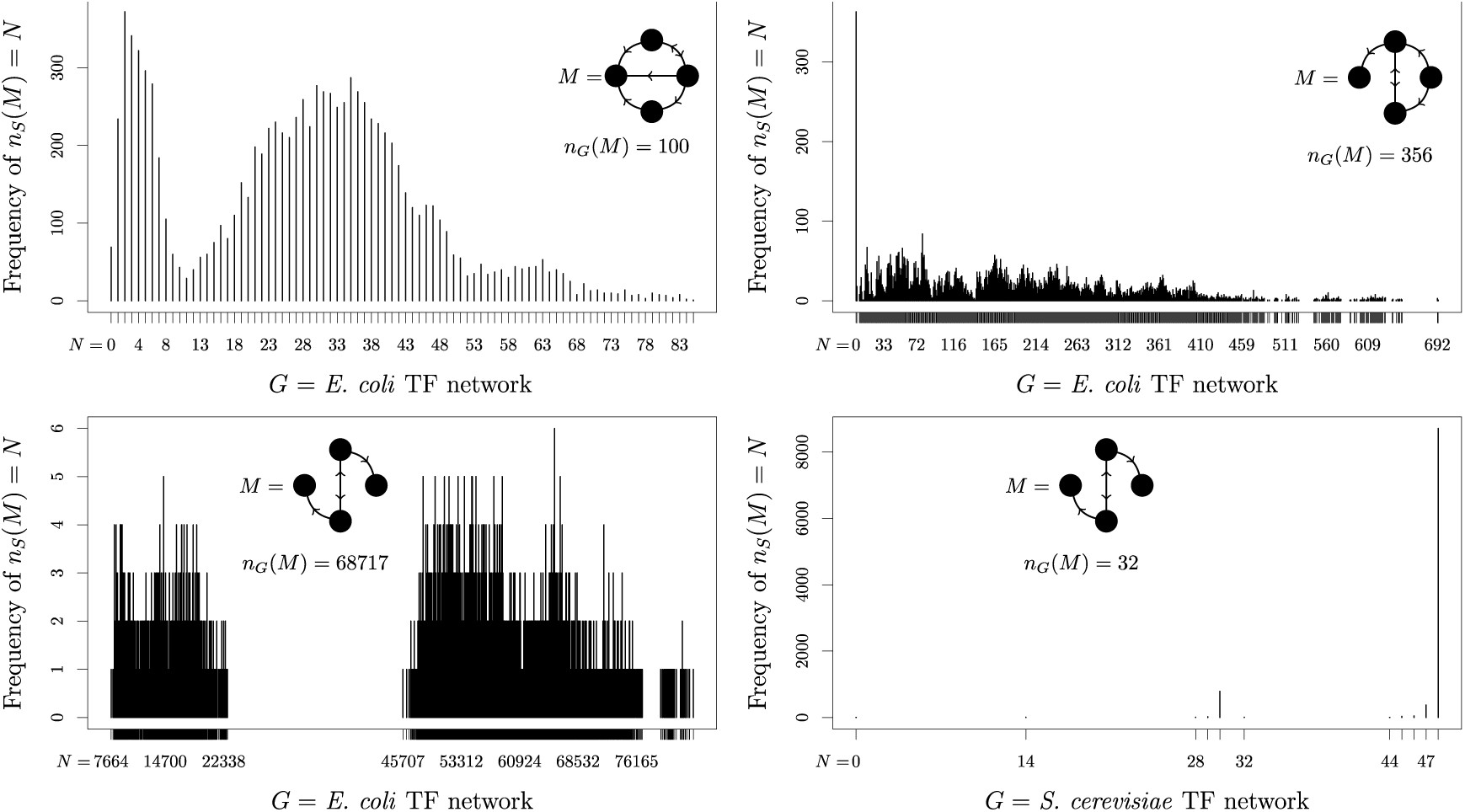
Example frequency distributions of the number of copies *n_s_*(*M*) of the candidate motif *M* found in networks similar to four bacterial transcription factor networks, as approximated by drawing 10,000 comparison networks from each 𝕊(*G*). The actual frequency *n*_*G*_(*M*) of the candidate motif *M* in the original network is also shown for comparison.

In some cases, instead of computing *p*-values, Z-scores are used directly for significance testing [21]. For example, Kashtan et al. argued that “for large networks and subgraphs, a high cutoff of Z = 5 or 10 can be used to detect significance using the sampling algorithm. Setting the Z-score cutoff to high values is important also for avoiding false positives” [8]. While not explicitly requiring that motifs are normally distributed, this approach amounts to an implicit assumption of normality, because if the underlying distribution is not approximately normally distributed then there is no guarantee that high Z scores correspond to especially unlikely outcomes. Such misapplication of Z-score tests can give rise to grossly inaccurate results. For example, Vinayagram et al. report some extremely high Z-scores of up to 2011.9, which assuming a normal distribution corresponds to a *p*-value of approximately 10^−878959^. However, using a crude lower bound for *p* (see SI Section S2.1 for full details), any motif from the network in question must have p ≥ 10^−20538^. The implied *p*-value therefore falls below this lower bound by at least 858,421 orders of magnitude, indicating that for this example the implicit assumption that candidate motifs follow a normal distribution cannot possibly hold. In such cases where the normality assumption fails to hold, Z-scores cannot be used to evaluate statistical significance.

In addition to not always following a normal distribution, frequency distributions of candidate motifs are not necessarily even reproducible over repeated runs. The switching method algorithm, which forms the basis for many of the most common software packages used in biological research (including FANMOD) to generate sets of similar networks, is known not to sample uniformly or independently [22], but is still commonly used because it is fast. The implicit assumption is that if sufficiently many similar networks are sampled from 𝕊(*G*), then these drawbacks have little practical effect. We show, however, that this is not always the case, as four separate runs of the switching method on the *E. coli* TF network each produce distinctly different results (Figure 3). Not only are the four histograms dissimilar, the estimated *p*-value for the indicated motif candidate ranges from 0.011 (not significant) to *10*^-29^ (highly significant). Such widely divergent results from one run to the next render any reported *p*-values in such examples effectively impossible to interpret.

**Figure 3:**
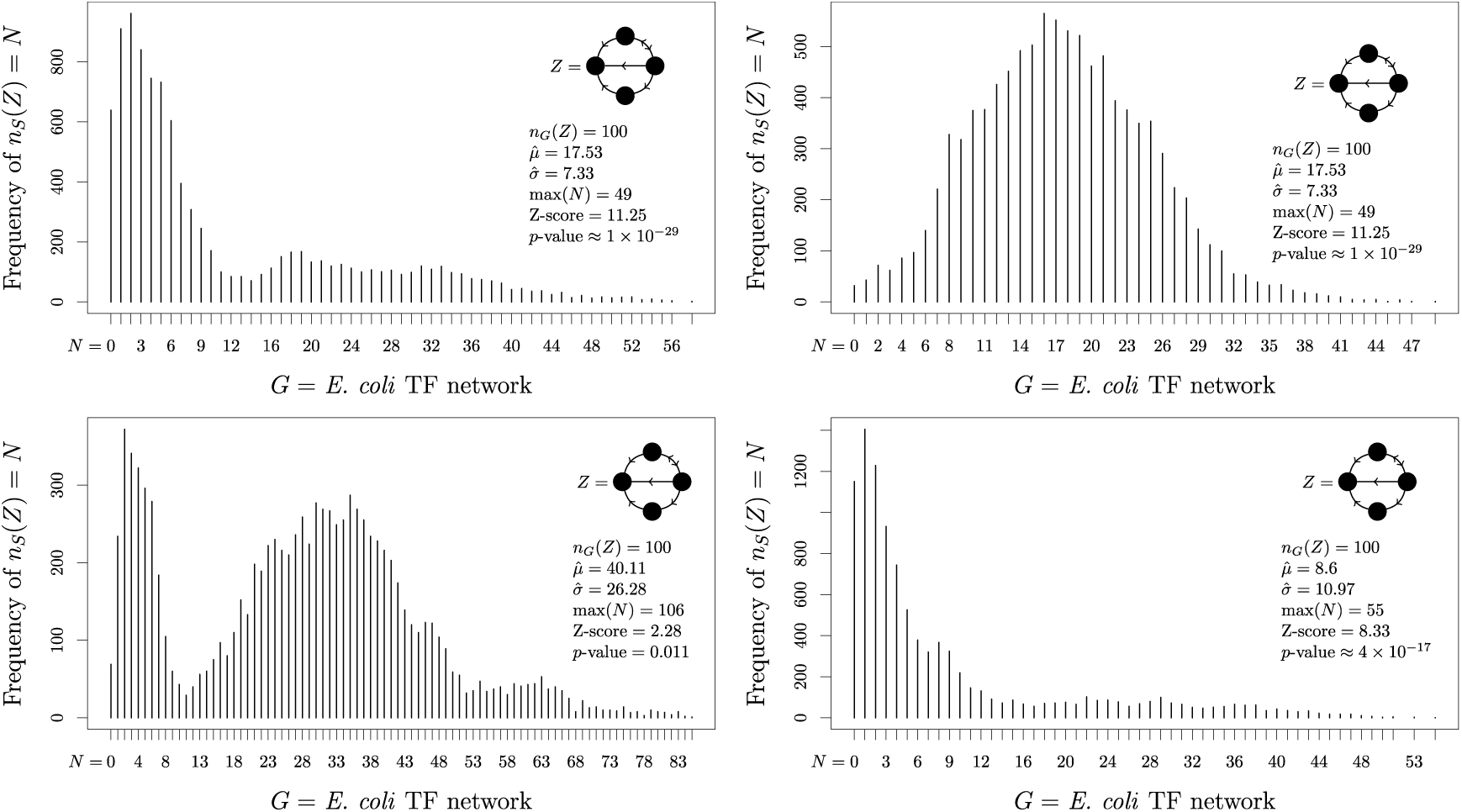
Frequency distributions of the number of copies *n*_*s*_(*Z*) of the motif-candidate *Z* found in networks similar to the *E. coli* TF network, as approximated by drawing 10,000 comparison networks from *𝕊(G*), generated using the switching method. Each of the four histograms is computed from a separate run of the algorithm, using the same candidate motif and the same input network. The actual frequency *n*_*G*_(*Z*) of the candidate motif *Z* in the original network is also shown for comparison.

## The independence assumption

A persistent problem with multiple hypothesis testing is that hypotheses are tested individually, not jointly, with the same search procedure simply repeated for each possible motif candidate [8, 19]. Such independent testing of multiple hypotheses is only valid under the assumption that different motif candidates are distributed independently in the underlying family of similar networks 𝕊(*G*). We show, however, that there are many circumstances in which this assumption fails to hold, and the occurrence of one motif is correlated with one or more others. To illustrate such correlation in a synthetic example, consider a network which contains three types of nodes: ‘source nodes’, with one outgoing edge and no other types of edges; ‘sink nodes’, with one incoming edge and no other types of edges; and ‘pipe nodes’, with exactly one incoming and one outgoing edge and no bidirectional edges. The class of networks considered here has *a* source nodes, *b* pipe nodes linking the source nodes to the sink nodes, and *a* sink nodes. Cyclic paths (loops) can occur among sets of pipe nodes, but not among source or sink nodes. An example of one such network is shown in Figure 4. In this example we consider the putative motif structure *L*_3_, which is simply a 3-edge-long path.

**Figure 4:**
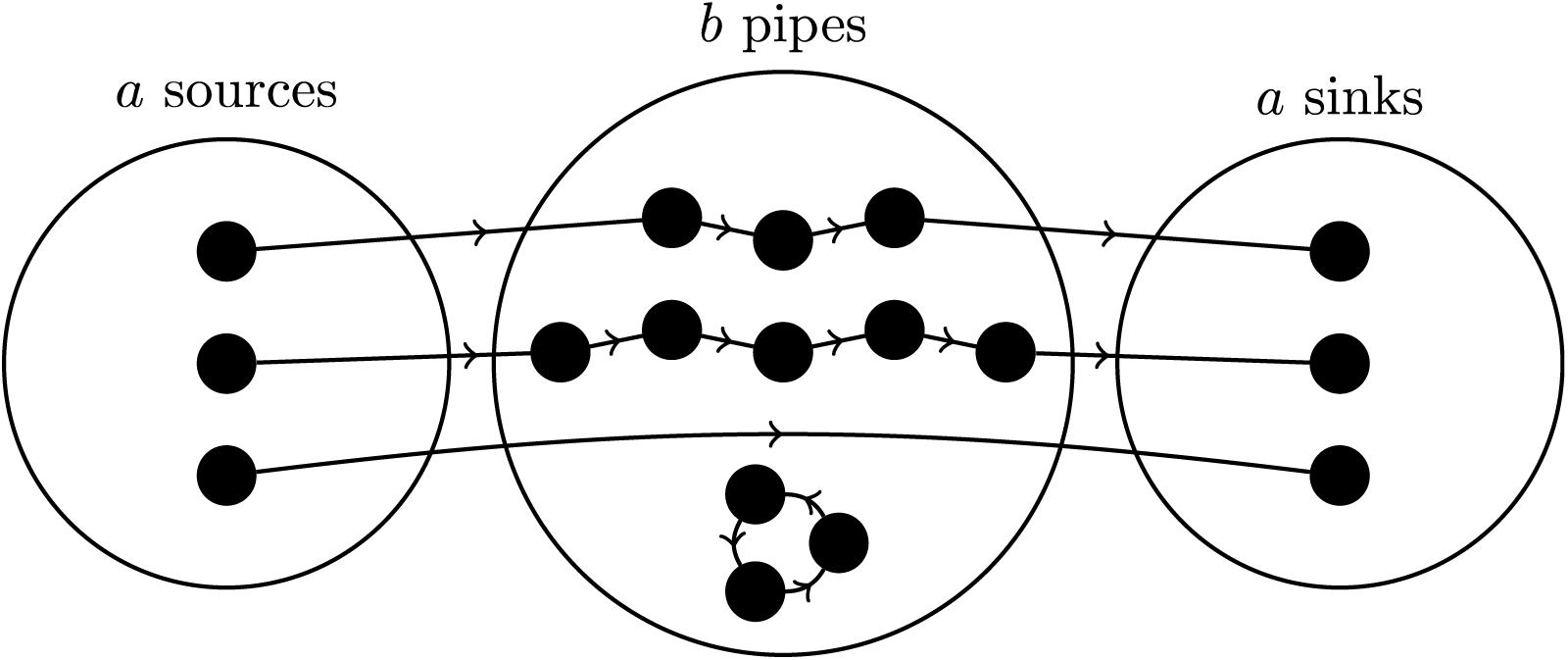
One member of a class of networks which exhibits correlation between the occurrence of different motifs.

The paths found in this network can be divided into three classes: non-cyclic (NC) paths which link source to sink via some number of pipe nodes, cyclic (C) paths which do not involve source or sink nodes, and source-sink (S) paths in which a source node connects directly to a sink without passing through any intermediate pipe nodes. We use *s* to denote the number of S-paths. Each NC-path of length *l* contributes *l − 1* copies of the *L*_*3*_ motif, while each C-path of length *l* (denoted *C*_l_) contributes *I* copies of the same motif as long as *l > 4* (and zero otherwise). We also know there are *a — s* NC-paths: one for each of the *a* source nodes, minus one for each of the *s* S-paths. Hence, the total number of copies of the *L*_*3*_ motif in networks of this form is:

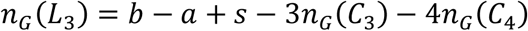

Given the linear relationship between *n*_*G*_(*L*_*3*_), *n*_*G*_(*C*_*3*_), and *n*_*G*_(*C*_*4*_), it is clear that the number of occurrences of candidate motifs *L*_*3*_, *C*_*3*_, and *C*_*4*_ in this type of graph is far from independent of one another. Such correlations are commonplace in real-world data, including the *S. cerevisiae* TF network (Figure 5a) and also the *E. coli* TF network (Figure 5b).

**Figure 5:**
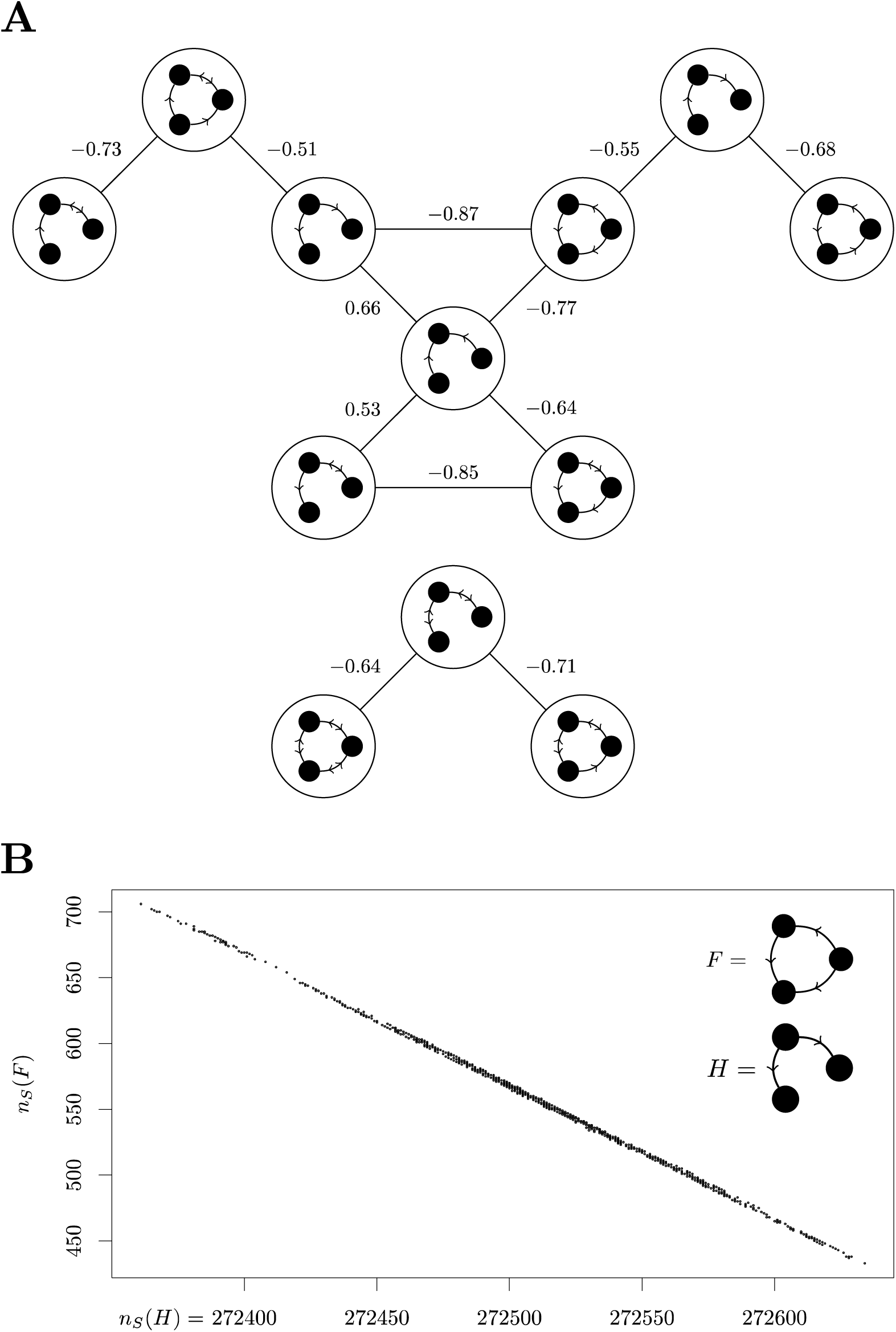
A A diagram depicting the Pearson correlation coefficients between *n*_*s*_(H) and *n*_*s*_(*H′*) for pairs of 3-node subgraphs *H* and *H′*. The input network is the *S. cerevisiae* TF network, and only coefficients of magnitude greater than 0.5 are shown. B: An example of correlation in the frequency of subgraphs *F* and *H* in the set 𝕊(*G*) of graphs similar to the *E. coli* TF network, with a Pearson correlation coefficient of −0.999.

While it has been argued that such correlations are “artifacts of the algorithm used to generate the ensemble of randomised networks” [23], we show theoretically that correlations occur even with uniform sampling. Whenever such correlations occur, methods which attempt to test for the existence of a wide range of different motifs without taking correlations into account will fail to deliver accurate results.

## Frequency vs. concentration

Thus far we have discussed *p*-values computed on the basis of the *frequency* of occurrences of subgraphs in a network. Many available network motif detection packages, however, instead compare subgraphs based on their *concentration* in a network [19]. Concentration is calculated as the number of occurrences *n*_*G*_(*H*) divided by the total number of connected subgraphs with the same number of nodes. In many cases in the literature little distinction is made between these methods, with the choice of which to use largely determined by computational convenience rather than theoretical considerations. In Figure 6, however, we present a number of cases from the *E. coli* metabolic network in which frequency and concentration statistics differ dramatically, illustrating that the choice of which method to use can drastically affect the results obtained.

**Figure 6:**
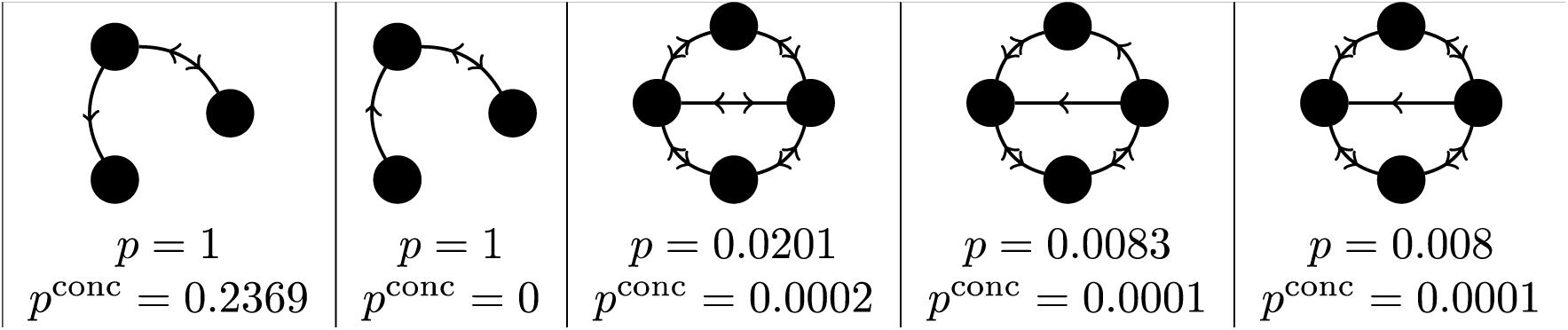
Some examples of putative motifs from the *E. coli* metabolic network where the computed frequency *p*-value *p* and computed concentration *p*-value *p*^conc^ differ considerably.

For a synthetic example in which frequency and concentration statistics are contradictory, we consider a network *G* with *k* nodes that have exactly one outgoing and no other types of edges and 3*k* nodes that have exactly 1 incoming edge, possibly one additional bidirectional edge, and no outgoing edges. In total, the network will have *m* bidirectional edges, with *m 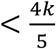*. One example of such a network is given in Figure 7. We show that for any two graphs selected from the set 𝕊(*G*), the concentration of the subgraph *T* (shown in Figure 7) is higher if and only if the number of occurrences of *T* is lower (see SI Section S2.5, for full details). Thus, *T* can be a motif according to its *p*-value only if it is an anti-motif (meaning that it occurs *less* often than expected in the set 𝕊(*G*)) according to its *p*^*conc*^-value, and vice versa. Which type of *p*-value to use is therefore not merely a matter of computational convenience, but can have substantive effects on the outcome of motif analysis.

**Figure 7:**
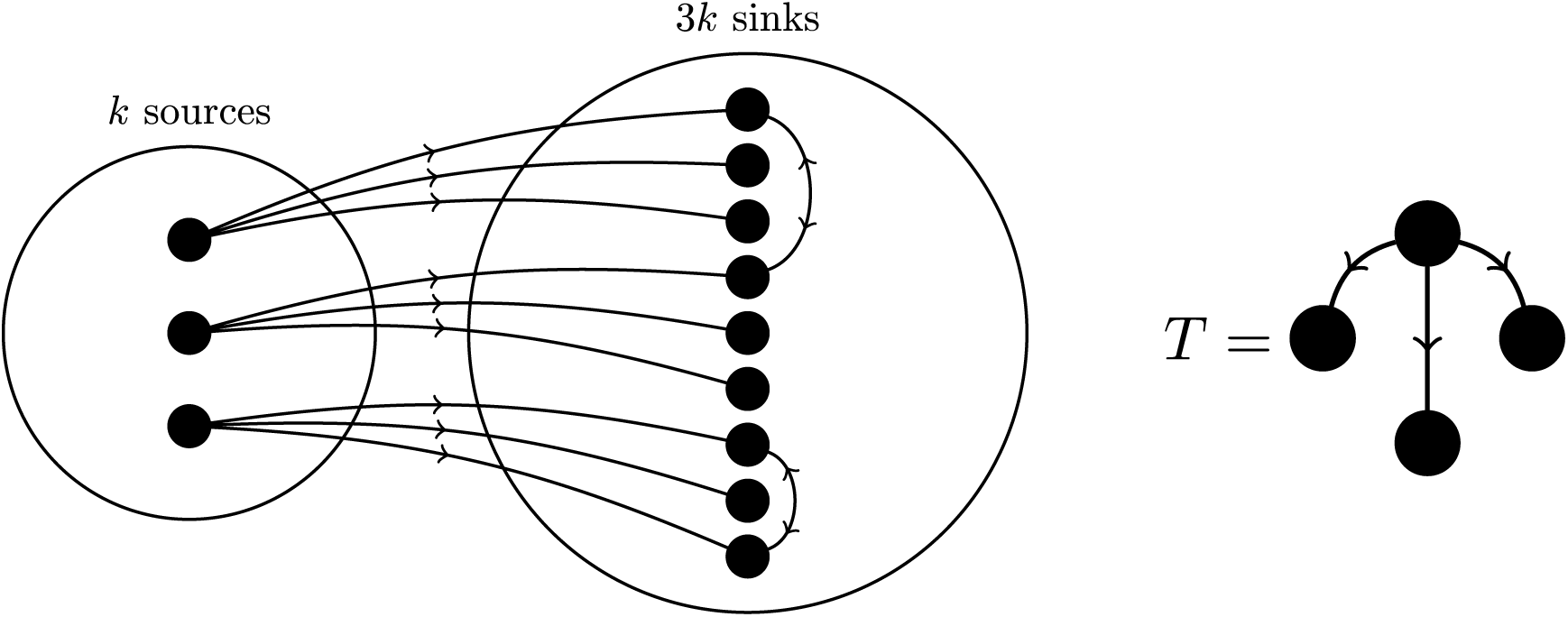
In this network, the subgraph *T* can be a concentration-based motif only if it is a frequency-based anti-motif, and vice versa.

## Conclusion

The ability to reliably identify network motifs is critical for understanding the complex regulatory networks that govern gene transcription and regulation, particularly in research concerning the prediction, organisation, and linking of transcription factors, binding sites, and promoters. Network motif identification can aid in overcoming the critical genomics bottleneck of processing increasingly enormous amounts of newly generated data. In addition, network motifs will play a pivotal role in the rational design and engineering of complex biological systems underpinning the field of synthetic biology. Given its importance, it is crucial that methods of identifying network motifs are statistically robust. However, we have demonstrated both theoretically and empirically that a collection of implicit assumptions present in mainstream network motif studies cannot always be assumed to hold, with the ramification that results from motif studies are not as mathematically unambiguous as is often presented. In particular, we showed that the assumption of normality does not always hold, and that non-normality results in materially inaccurate *p*-values, invalidating the standard methods of inferring that a given network structure is present at above-chance levels and thereby constitutes a true motif. We showed that the frequencies of different motif candidates are not always independent of one another, and that such lack of independence invalidates methods which test for the presence of large numbers of motifs independently of one another. Finally, we showed that motif frequency and motif concentration are not interchangeable, and that the choice of which to use can have substantial effects on the outcome of any analysis. We conclude that it is unclear whether current mainstream network motif identification methods have the capacity to reliably differentiate between spurious results and events of true statistical significance - a basic requirement for a mass-hypothesis testing tool.

## Methods

Condensed descriptions of methods are described below; see SI Methods for details. We analysed three real-world networks; the *E. coli* TF network (sourced from RegulonDB [24]), the *S. cerevisiae* TF network (sourced from [25]) and the *E. coli* metabolic network (we use the version packaged with Kavosh [26], where it is attributed to Kyoto Encyclopedia of Genes and Genomes (KEGG) [27]). Both organisms are the best known and characterized genome model organisms. We take these networks ‘as is’ from their respective sources, without introducing additional (and potentially arbitrary) edge thresholding criteria or other post-process manipulations. For each of these three networks, a set of 10,000 similar networks 𝕊(*G*) was generated using the software NetMODE, which uses the switching method (see SI Section S4.4.2 for more information). The number of occurrences of the motif candidate of interest was then calculated using the same software. Correlations between motif candidates, *p*-values, and frequency histograms were computed and plotted using R scripts.

## Acknowledgements

AMB acknowledges support from the National Health and Medical Research Council (1022688). RJS was supported by her NSFC Research Fellowship for International Young Scientists (grant numbers: 11450110409, 11550110491), NSF China grant 61170301, and the Thousand Youth Talents Plan in Tianjin.

## Author Contributions

RJS and MB designed the study and performed the analysis. JF, RJS, MB and AMB wrote the manuscript.

## Author Information

The authors declare no competing financial interests.

Correspondence and requests for materials should be addressed to James Fodor and Ashley Buckle.

